# In Vivo Flow Cytometry of Extremely Rare Circulating Cells

**DOI:** 10.1101/488619

**Authors:** Xuefei Tan, Roshani Patil, Peter Bartosik, Judith Runnels, Charles P. Lin, Mark Niedre

## Abstract

Circulating tumor cells (CTCs) are of great interest in cancer research, but methods for their enumeration remain far from optimal. We developed a new small animal research tool called “Diffuse *in vivo* Flow Cytometry” (DiFC) for detecting extremely rare fluorescently-labeled circulating cells directly in the bloodstream. The technique exploits near-infrared diffuse photons to detect and count cells flowing in large superficial arteries and veins without drawing blood samples. DiFC uses custom-designed, dual fiber optic probes that are placed in contact with the skin surface approximately above a major vascular bundle. In combination with a novel signal processing, algorithm DiFC allows counting of individual cells moving in arterial or venous directions, as well as measurement of their speed and depth. We show that DiFC allows sampling of the entire circulating blood volume of a mouse in under 10 minutes, while maintaining a false alarm rate of 0.014 per minute. Hence, the unique capabilities of DiFC are highly suited to biological applications involving very rare cell types such as the study of hematogenic cancer metastasis.

## Introduction

There are many biological processes involving rare cells that circulate in the peripheral blood. Circulating tumor cells (CTCs) are extremely rare (fewer than 100 cells per mL) and are critical in the development of cancer metastasis. Numerous studies have shown that CTC burden correlates with disease progression and response to treatment^1,2^. Existing methods for enumeration of CTCs remain far from optimal. Normally, blood samples are drawn and target cell populations are isolated using a variety of *in vitro* assays such as flow cytometry, size-based separation, immuno-magnetic separation, and microfluidic capture^3^. These have provided a wealth of information, for example in combination with fluorescence imaging or high-throughput sequencing methods. However, it is well established that the general methodology of drawing, enriching and analyzing blood samples may be problematic^4,5^. Blood samples are known to degrade rapidly after removal from the body^6^. Moreover, analysis of small blood volumes (relative to the total peripheral blood volume) can lead to significant under-or over-estimation of CTC burden, simply due to sampling statistics^7,8^. In mouse studies, blood collection is limited to about 10% of the blood volume every two weeks without fluid replacement^9^, making longitudinal studies of CTC burden extremely difficult. The process of drawing blood can also trigger a stress response in the animal^10^.

To overcome this, researchers have developed ‘*in vivo* flow cytometry’ (IVFC) methods to count cells directly in mice without having to draw blood samples. Typically, these are modified intravital fluorescence microscopes that operate in trans-illumination mode through a mouse ear, and optically sample blood flowing through a small arteriole^5,11–15^. Photoacoustic IVFC methods have also been described^4,16^, which generally rely on detection of naturally highly-pigmented cells such as melanoma. In microscopy-IVFC, the circulating blood volume sampled is about 1 μL per min, so that it is normally used for applications involving thousands of circulating cells per mL^5^. For circulating cells at lower concentrations, mice usually must be euthanized and the entire peripheral blood volume drawn and analyzed. In this case, experiments are terminal and individual animals cannot be followed longitudinally over time to track disease progression.

In recent years, our team has explored the use of highly-scattered (‘diffuse’) light for studying rare fluorescently-labeled circulating cell populations in bulk tissue^17–19^. The rationale is to interrogate major blood vessels where flow rates are orders-of-magnitude higher than in an arteriole of the ear, where intravital microscopy methods are not readily applicable. We recently described a fiber-optic probe design composed of a set of bundled source and detection fibers, with integrated interference filters and collection lens deposited directly on the tip. We showed that when placed on the surface of the skin above a large blood vessel, this probe allowed sensitive fluorescence detection of cells moving in bulk tissue. The probe had a number of advantages including, simple alignment, minimal artifacts due to limb movement and ready translation to larger limbs and species. However, there were also a number of limitations which precluded its use in a real cancer metastasis studies, in particular, i) it could not distinguish cells moving in arterial, venous, or capillary bed, making it susceptible to over-counting of cells on their return trip through the vasculature, ii) it was susceptible to false-positive signals due to electronic noise or motion, and, iii) the measured count rate was lower than predicted based on the cell concentration, which we later determined was primarily due to hypothermia and poor blood flow in the limb while the mouse was under anesthesia.

In this paper, we report the design, validation and in vivo characterization of a new instrument called ‘Diffuse *in vivo* Flow Cytometry’ (DiFC). DiFC builds on our previous work, but introduces a number of critical advances which, in combination allowed us to sample hundreds of microliters of blood per minute while maintaining a negligible false alarm rate. First, we used two improved fiber bundle probes, and developed a pairwise coincidence detection algorithm for ‘matching’ detected cells between the optodes. As we show, this allowed us to distinguish cells moving in arterial or venous (forward and reverse direction) flow, and calculate the cell speed and depth. It also allowed us to virtually eliminate false positive signals, and discard signals from cells moving in the capillary bed. Second, we achieved better thermal control of the mouse tail while under anesthetic, avoiding hypothermia in the limbs and significantly increasing blood perfusion in the sampling volume versus our previous work.

We first tested and characterized DiFC with multiple myeloma (MM) cells labeled with a near-infrared fluorescent dye. We demonstrated that it allowed sampling of 284 μL of peripheral blood per minute, so that the entire ~2 mL circulating blood volume of a mouse could be sampled in under 10 minutes. Use of the matching algorithm allowed us to aggressively reject false positive signals due to electronic noise or motion, and as we show yielded a low false alarm rate (FAR) of 0.014 per minute. Moreover, DiFC operates continuously and non-invasively, so that it can monitor cell population kinetics longitudinally over time in the same animal. The optical design of DiFC is such that it can be used with a wide variety of fluorophores and fluorescent protein-labeled cell lines^5^. Finally, because DiFC works in epi-illumination and detection geometry (surface contact versus trans-illumination), is also advantageous in that it is readily scalable to larger limbs and species.

### DiFC Instrument

A schematic of the DiFC instrument is shown in ***Figure 1a***. DiFC uses two custom designed fiber-optic probes separated by 3mm (center-to-center). The individual probe was similar to the prototype design we described previously^18^, although a number of components in the optical system were upgraded. Each probe consisted of a central excitation fiber that delivered laser light to the skin surface, and eight fibers for collection of near-infrared fluorescent light. The excitation source was an upgraded 642 nm laser which provided 20 mW at the sample per probe, thereby improving the detection signal-to-noise ratio (SNR) compared to our prior work. Fluorescent light generated by single moving cells is extremely weak relative to background auto-fluorescence, so that careful optical design to efficiently collect cell fluorescence, and minimize laser-induced auto-fluorescence generation in the fibers was critical. Each probe had a micro-machined 640 nm bandpass and 660 nm longpass filter mounted directly on the tip. A miniaturized aspheric lens also improved collection of fluorescent light (***Fig. 1b***). On the detection side, the collection fibers were split into two bundles that terminated on photomultiplier tubes (PMTs) filtered with 700 nm bandpass filters. In this work, the detection filters were at the same wavelength, and the signals for the 2 PMTs were summed, which further improved the SN). In the future, we could also use different filters on the PMTs to allow multiplexed detection of cells labeled with different fluorophores. The PMT outputs were amplified with low noise current-preamplifiers and then acquired with an analog-to-digital converter. Specific optical components for are listed in detail in the ***Methods*** section below.

**Figure 1.**
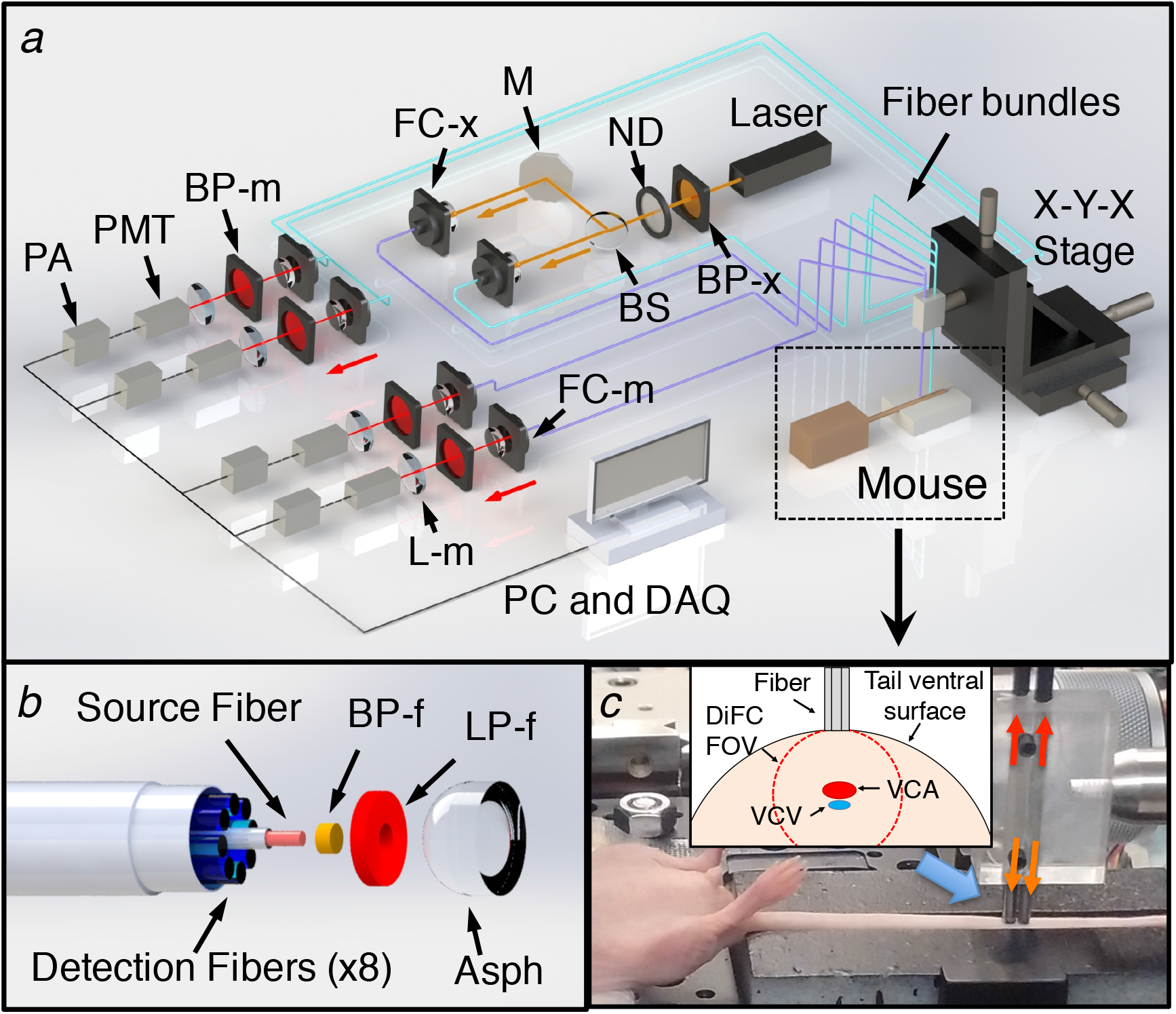
(a) Schematic of the DiFC instrument. See text for details. (b) DiFC fiber-optic probes contain a source fiber, 8 collection fibers, and integrated filters and collection optics. The probes were mounted in a holder (c) and placed on the ventral skin surface of the mouse tail. (inset) stylized cross-sectional view of the tail and DiFC FOV. The ventral caudal artery (VCA) and ventral caudal vein (VCV) are approximately 1 mm deep.

Mice were held under inhaled isofluorane during DiFC scanning to prevent movement and were kept warm using two warming pads. One was placed under the body, and a second one was place over the exposed area of the tail. The latter was found to greatly improve blood flow, and as we show increase the linear flow speed of cells in the artery significantly. The probes were placed in firm contact with the skin surface, approximately over the large ventral caudal (VC) bundle in the mouse tail (***Fig. 1c***). Alignment over the blood vessel was simple and was achieved visually and by translating the probe tip on a manual X-Y-Z translation stage.

### DiFC Data Analysis

***Figures 2a and b*** show example data traces measured from the ventral surface of the tail of a PBS-injected control mouse (***Fig. 2a***) and a mouse injected intravenously with multiple myeloma (MM) cells labeled with the near-infrared cell trace far red (CTFR) dye (***Fig. 2b***). As fluorescently-labeled cells passed through the DiFC field-of-view (FOV), fluorescence ‘peaks’ were measured, similar to our previous work^18^. We first performed basic pre-processing on the data including subtraction of the approximately static auto-fluorescence background (~10 μA) and application of a 3-point (3ms) moving average filter to reduce noise.

**Figure 2.**
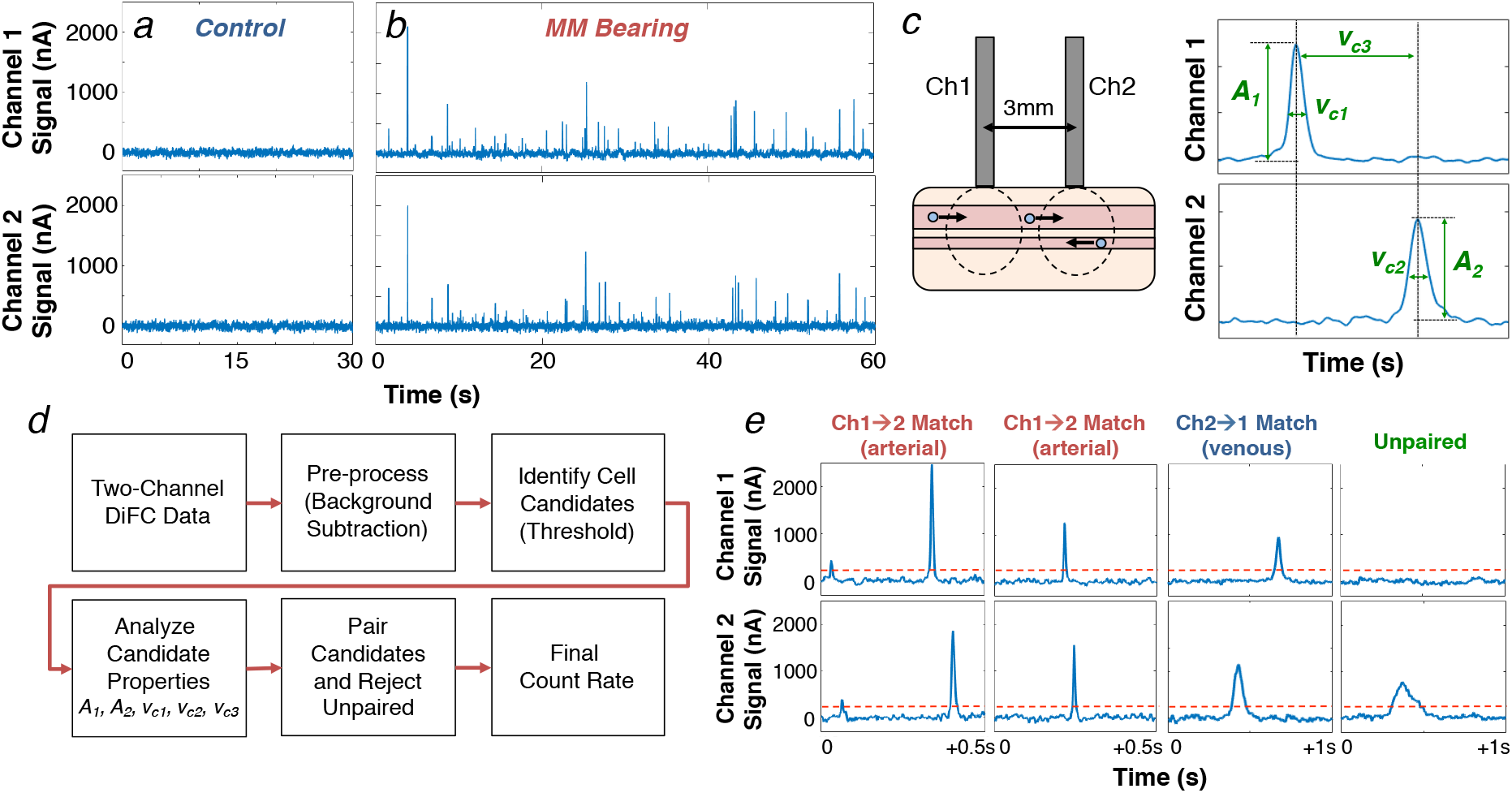
Representative data take with DiFC from (a) sham PBS-injected control mouse, and (b) CTFR-labeled MM.1S injected mouse. The peaks indicate cells moving through the instrument field of view. (c) We extracted the amplitude, width and estimated linear speed of detected peaks, given the known fiber-separation. (d) we developed an algorithm to determine the direction of cell travel, and remove unpaired peaks (see text). (e) Examples of forward, reverse, and un-paired peaks are shown.

Monte Carlo analysis of the DiFC FOV indicated that the instrument collects light from the first 2 mm of tissue. When placed on the ventral surface of the mouse tail (***Fig. 1c inset***), fluorescent light is therefore dominantly collected from cells in the ventral caudal bundle – the largest vessel of which is the ventral caudal artery (VCA)^20^. However, it can also originate from any number of smaller blood vessels including the ventral caudal vein (VCV) or the capillary bed. Occasionally, electronic noise or motion artifacts can also result in false positive ‘peaks’ in the signal.

To address this, and also accurately determine the speed and depth of cells, we developed an algorithm to jointly use the data from the two probes (see ***Methods*** for details). Briefly, ‘candidate’ peaks on were first identified by comparison to a fixed threshold. Candidates on both channels were then analyzed for their amplitude (***A_1_, A_2_***), speed estimated from the spike widths (***v_c1_, v_c2_***), and speed estimated from the time-separation between channels (***v_c3_***) (***Fig. 2c***). Peaks were then paired according to amplitude and speed similarity in either the forward (channel 1-to-2) or reverse (channel 2-to-1) directions. Cell candidates that were un-paired or coincidental (implying an artifact) were rejected.

The overall approach is summarized schematically in ***Fig. 2d***. Example measured signals shown in ***Fig. 2e***, where paired cells moving in the forward and reverse directions are shown, as well as an example of an unpaired signal that is rejected by the algorithm.

### Detection and Counting of MM Cells *in Vivo* with DiFC

We tested DiFC by tail vein (i.v.) injection of either 2.5 x 10^4^ or 10^5^ CTFR labeled MM cells (N = 5 each) in a 200 μL bolus in nude mice. DiFC scanning was performed starting 10 minutes after injection for approximately 90 minutes for each mouse. One advantage of DiFC is that the cell count rate can be measured continuously while the mouse is under anesthesia, so that circulating cell populations that change over minutes or hours can be measured. An example DiFC data trace from a mouse injected with 10^5^ CTFR-labeled MM cells is shown in ***Figure 3***. Only the data from ‘channel 1’ is shown here for clarity, but data was collected from both channels continuously, which allowed the algorithm to match cells in the arterial, and venous directions. Unmatched peaks were presumed to be either cells moving in smaller blood vessels or motion artifacts, and were therefore discarded by the algorithm. Previous work by our group and others has shown that most cells clear in the first pass through the vasculature, and then continue to clear from circulation into the bone marrow niche^21^ as is observed here.

**Figure 3.**
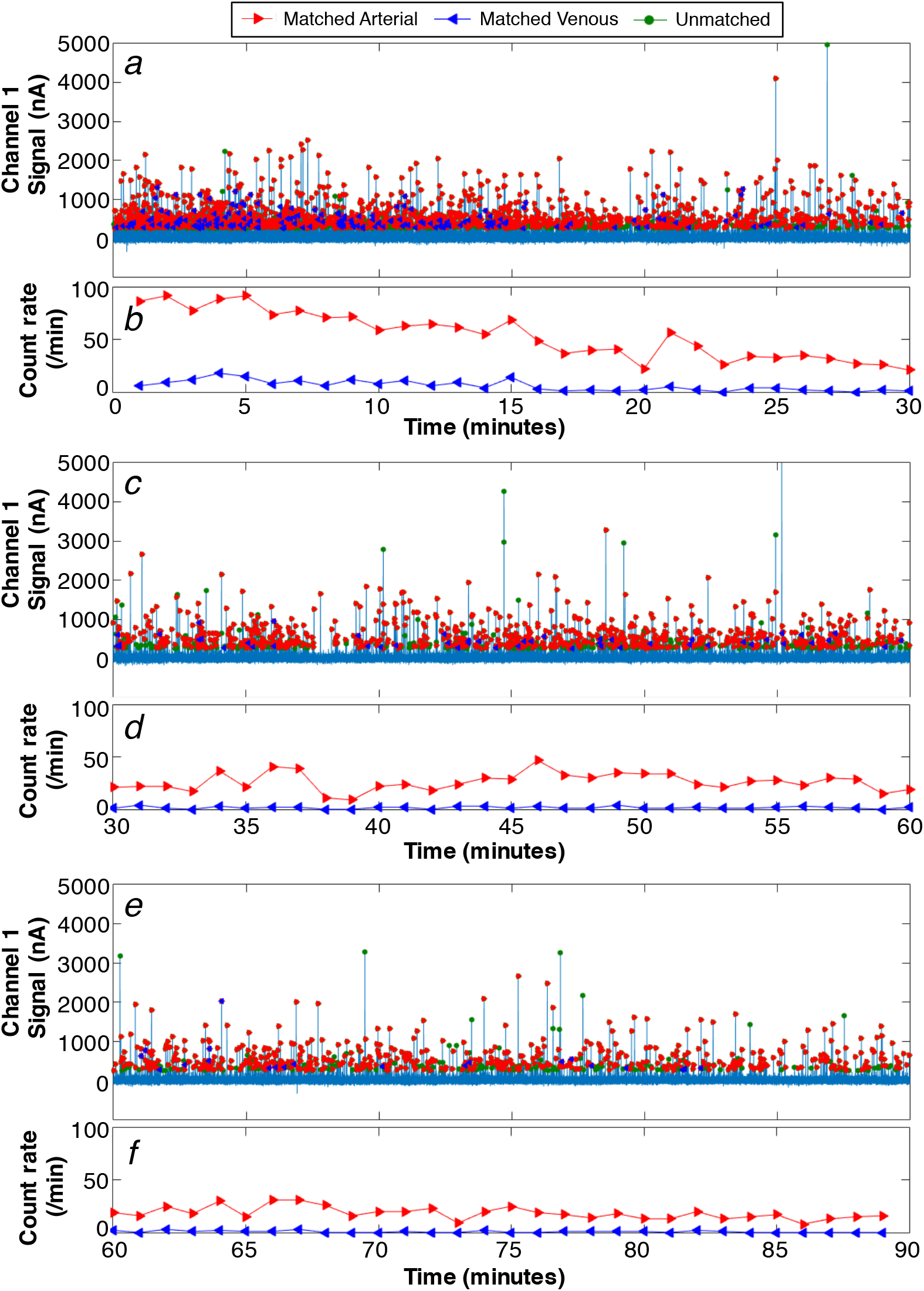
Example data collected from a mouse injected with 10^5^ CTFR labeled cells and 10^6^ unlabeled cells. (a) Processed DiFC data for 1 channel, showing identified arterial, venous and unmatched peaks in the first 30 minutes of the scan. (b) The count rate per minute in the arterial and venous directions, in the first 30 minutes. (c,d) corresponding data for 30-60 minutes and (e,f) 60-90 minutes of the scan are shown. The count rate declines over time as cells clear from circulation as expected.

We also performed sham (PBS) injections on an additional 7 control mice and acquired data for 60 minutes each. Example DiFC data measured from two control mice are shown in ***Figure 4***. For the second mouse (***figs. 4c,d***) a few individual false-positive ‘candidate peaks’ were detected on a single channel (green circles), but as discussed in more detail below these were rejected by the matching algorithm.

**Figure 4.**
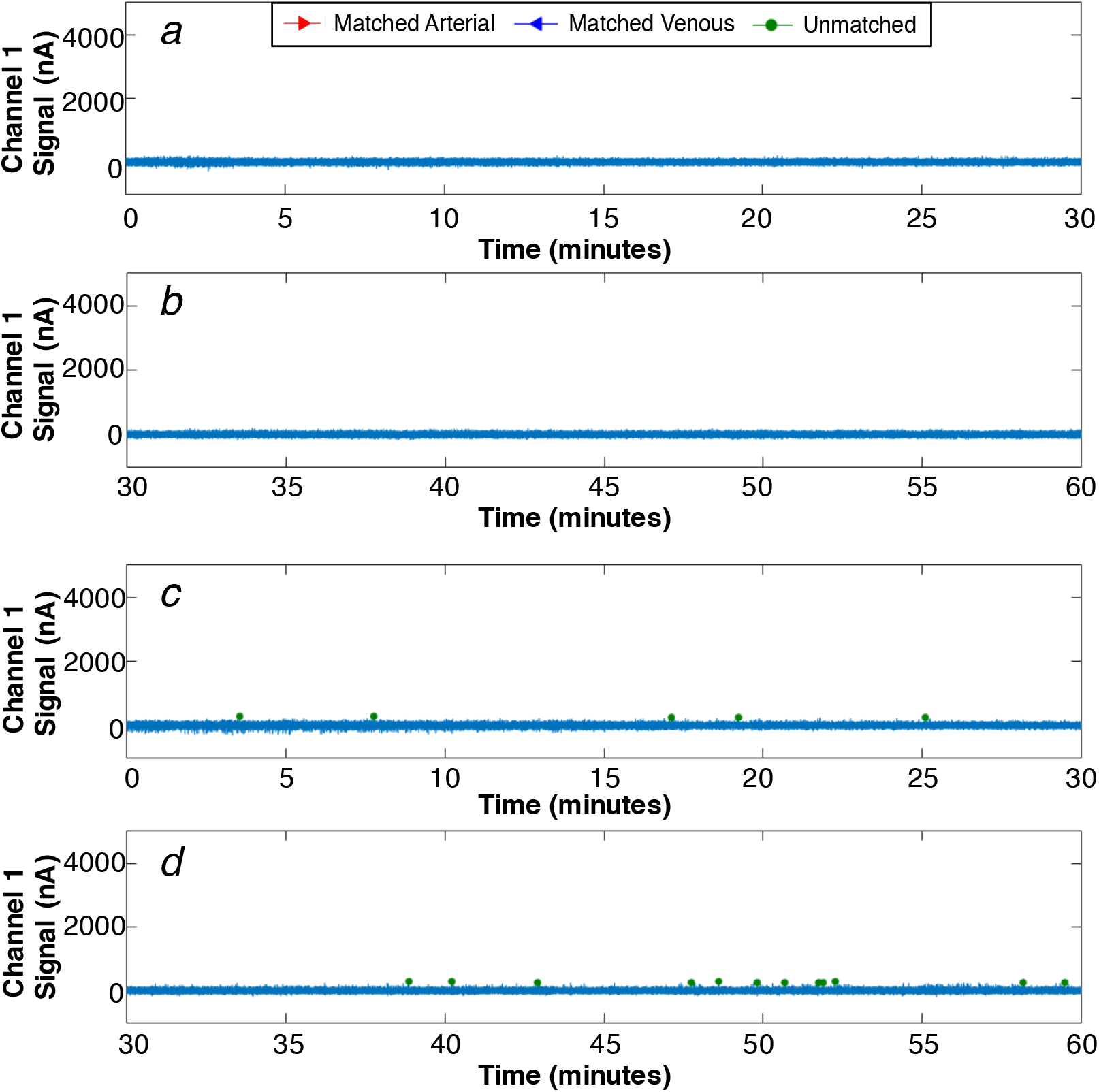
Example processed 1-channel DiFC data collected from a PBS-injected control mouse for (a) 0-30 minutes and (b) 30-60 minutes of the scan. (c,d) example data collected from a second example control mouse which was slightly noisier and produced a number of small false alarm peaks (green circles). However, the peak-matching algorithm eliminated these.

The data from all MM-bearing mice is summarized in ***figure 5. Fig. 5a*** corresponds to the data collected from a mouse injected with 10^5^ labeled cells (the same animal as ***figure 3)*** and shows the count rate for cell candidates on channels 1 and 2, along with the count rate for matched peaks in the forward (arterial) and reverse (venous) directions in 10-minute intervals.

**Figure 5.**
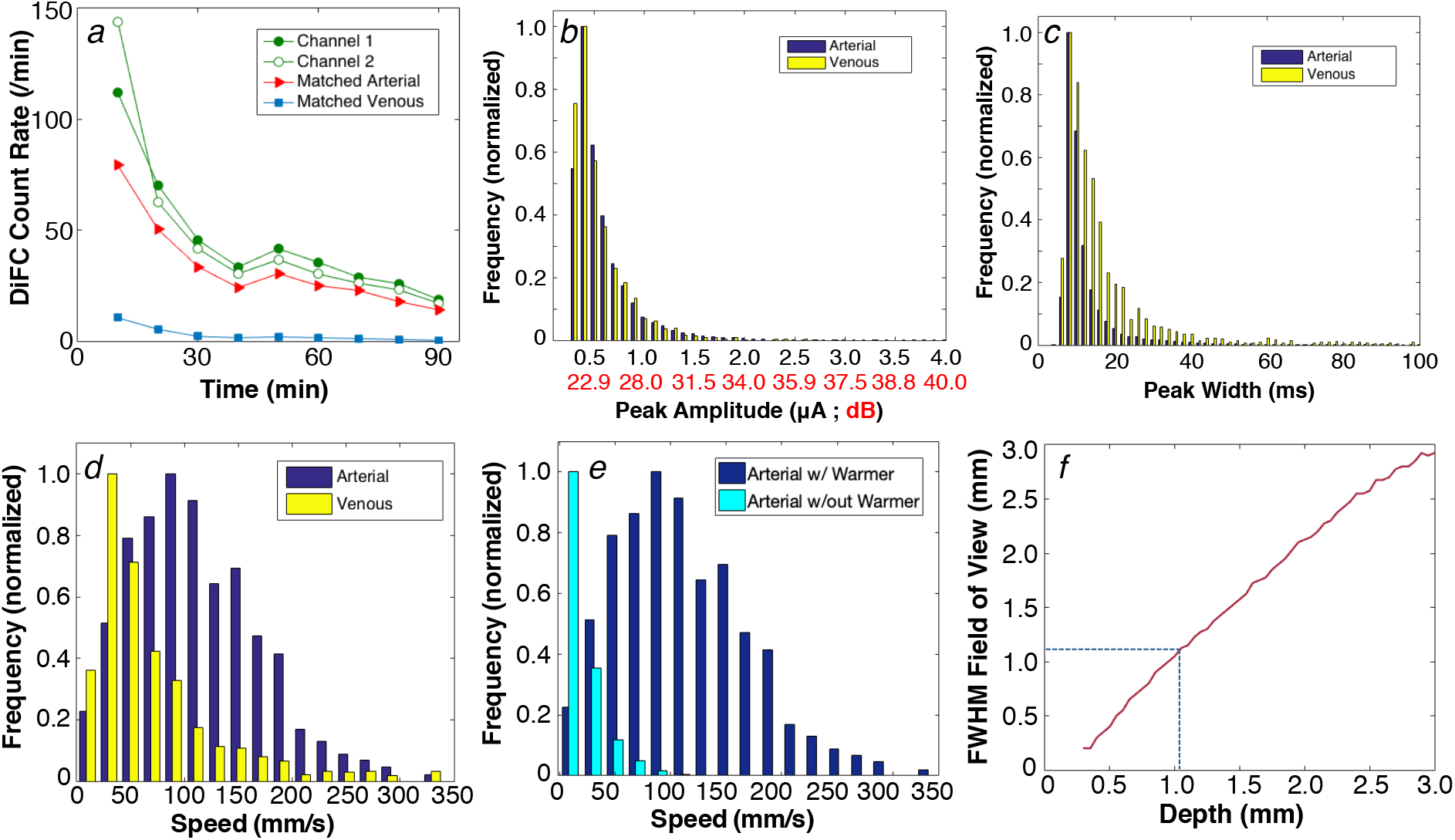
(a) Example data collected from an MM-injected mouse, up to 90 minutes following injection. The detected count rate for fibers 1 and 2, as well as the arterial and venous matched count rates are shown. The distribution of measured (b) peak amplitudes, expressed in PMT current (nA) and SNR (dB), (c) detected peak widths for arterial and venous matched cells, and (d) linear cell speeds for arterial and venous matched cells from all mice in this study are shown. (f) Use of a warming pad over the tail significantly increased the linear flow speed of cells moving in the artery. (g) The measured cell speeds and pulse-widths, in combination with a Monte Carlo simulation of the DiFC collection volume allowed us to estimate the depth of origin of the signals, which in this case was 1.1 mm (dotted line)

Overall, approximately 67% and 7.5% of cell candidates were matched in the forward (arterial) and reverse (venous) directions, respectively. The larger count rate in the artery is expected considering the larger size and flow rate of the VCA relative to the VCV. In terms of the detection signal-to-noise ratio (SNR) of detected peaks, the distribution peak amplitudes over all mice in arterial and venous directions is shown in ***Fig. 5b***. The mean peak heights were 532 and 503 nA in the arterial and venous directions, respectively. The average system noise *σ* was 40 nA from the 7 control mice tested, so that these peak amplitudes correspond to a SNR of 22.4 and 22.0 dB, respectively (*SNR = 20*log10(I/σ*), where *I* is the average peak amplitude). The similar SNR in both directions was expected since the VCA and VCV are at similar depths in the ventral caudal vascular bundle (with the vein being only slightly deeper than the artery).

The algorithm also allowed us to measure the fluorescence peak widths as cells were detected by DiFC. Here, the full-width-at-half-maximum (FWHM) of matched peaks in the arterial direction were on average narrower (11.5 ms) than in the venous direction (17.8 ms) (***fig. 5c***). The narrower peak width is related to the cell speed, which could be directly calculated by considering the fiber probe separation (3mm) and the time difference between detections on the two probes, as shown in ***fig. 5d***. The mean cell speed over all mice was 112.3 mm/s in the arterial direction, and 76.6 mm/s in the venous direction. These speeds are generally consistent with the arterial speeds reported in mice by others^22^.

However, these flow speeds are significantly higher than we reported in our previous work^18^, which was estimated solely from the FWHM of detected peaks. The difference is primarily due to the use of the warming pad placed over the tail, which maintained blood perfusion throughout the scan. For comparison, we measured the flow speed in an additional 3 MM-injected mice where no heating pad was used as shown in ***fig. 5e***. For these mice, the average arterial flow speed was only 19.7 mm/s. Our new matching algorithm also allowed measurement of the speed in specifically arterial flow only, whereas our previous method averaged the estimated speed from all detected cells, which included slower venous flow as well as smaller blood vessels.

It was further possible to calculate the depth of the matched moving cells as follows: We multiplied the average peak widths in ***fig. 5c*** by the average cell speeds in ***fig. 5d*** and estimated that the DiFC field of view was 1.1-1.2 mm FWHM. Monte Carlo analysis of the DiFC probe sensitivity function in optically scattering tissue 12 shows that this width corresponds to an average tissue depth of approximately 1.1 mm^23^ (***fig. 5f***). This predicted depth agrees well with the approximate depth of the ventral caudal bundle (~ 1 mm) in an adult mouse^20^.

### DiFC False Alarm Rate

The matching algorithm also allowed us to virtually eliminate false-positive signals due to electronic or motion artifacts. These data are summarized in ***fig. 6a***. As shown, the average FAR from the matching algorithm was at least an order of magnitude lower than the FAR when data from a single fiber was used, regardless of the detection threshold. For CTFR-labeled MM cells, we used a counting threshold of 250 nA which was explicitly determined from the measured intensity of labeled cells compared to calibration microspheres on DiFC and conventional flow cytometry (see ***Method*s**). At this threshold, the FAR was 0.014 per minute, which is equivalent to one false alarm every 70 minutes. We also note that all of the false alarms came from a single animal (i.e. 6 out of 7 showed not false-alarms). Therefore, DiFC data is extremely stable over long acquisition periods.

**Figure 6.**
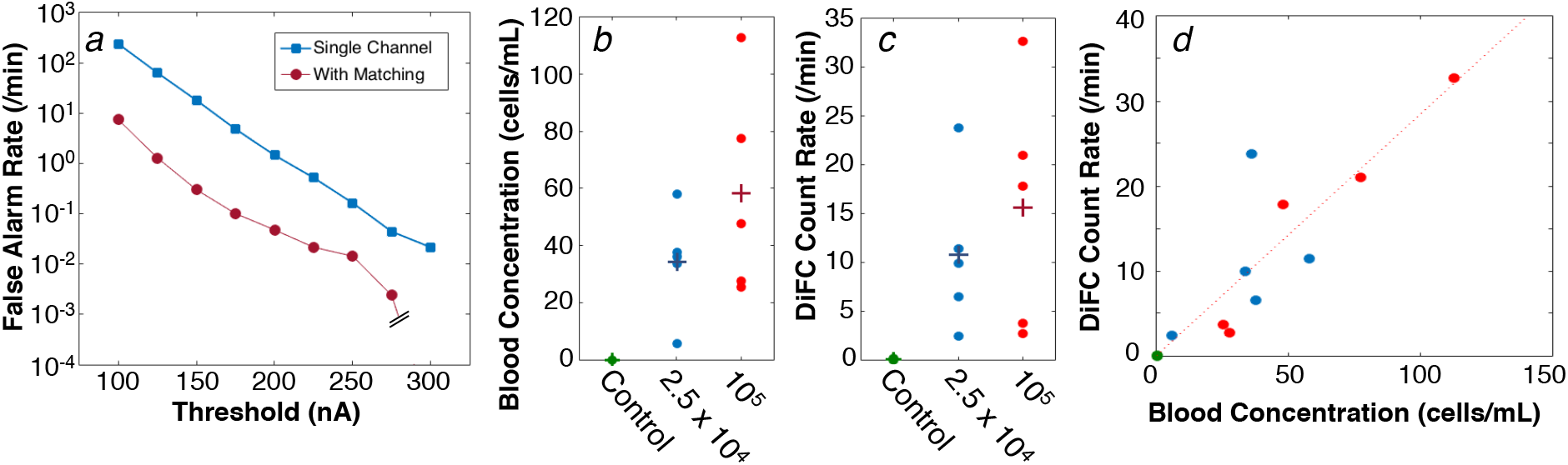
(a) False-alarm rate measured from sham-injected control mice for different detection thresholds. As shown, use of the matching algorithm drastically reduced false-alarm signals. At the operating threshold of 250 nA, the FAR was 0.014 per minute. (b) The number of cells in extracted blood was compared to the (c) DiFC arterial count rate during the last 10 minutes of scanning. (d) The result showed a good linear agreement between the count rate and MM cell burden.

### DiFC Detection Sensitivity

To determine the sensitivity of DiFC, we drew between 0.5-1 mL of blood from the mice after 90 minutes of scanning, and counted MM cells in the samples. Blood samples were diluted in heparin and PBS, and cells immediately counted *in vitro* using the DiFC system (see ***Methods***). We used this method since potential loss of cells in sample handling was a major concern,^5^ and this method required almost no handling or enrichment of the samples, which is known to induce cell loss. We took care to determine *in vitro* and *in vivo* cell counting thresholds relative to commercial calibration fluorescence microspheres so that counts from blood samples could be compared directly to DiFC data (see ***Methods*** and ***Supplemental Fig. S1***).

The number of MM cells per mL (CTC burden) for all MM-bearing mice (N = 10) and controls (N = 7) are shown in ***Figure 6b***. The low (< 150 cells/mL) concentrations measured were expected since it is known that *i.v*. injected MM cells clear rapidly from circulation, so that after 90 minutes only a few cells remain^18,21^. Likewise, the arterial DiFC count rate in the final 10 minutes of scanning before drawing blood is shown in ***fig. 6c***.

For both data sets, significant inter-experimental variability was observed, which we attribute to slight differences in injection efficiency and dilutions. However, when all data points were plotted together (***Fig. 6d***) a good linear relationship was observed with an r^2^-coefficient of 0.82. One outlier point was observed, which, when removed yielded an r^2^-coefficient of 0.93. It should also be noted that there is a relatively large uncertainty associated with the low blood cell concentration estimates Assuming Poisson counting statistics in ***fig. 6a*** and the x-axis of ***fig. 6d***, where the uncertainly for each point shown is 10%-25%.

Critically, the slope of ***Fig. 6d*** indicates that a concentration of 1 cell per mL produced one DiFC count every 3.52 minutes in the arterial direction. Equivalently this means that the volume of blood sampled with DiFC is 284 μL per min, indicating that the entire circulating blood volume of a mouse can be sampled in less than 10 minutes. This sampling rate is in excellent agreement with the estimated blood flow rate in the VCA of a mouse which we calculated as follows: the average measured cell speed in the artery was *v_s_* = 112.3 mm/s. The size of the VCA is unknown but we assumed it was a cylindrical with diameter of 250 μm^20^. Assuming a simple flow profile, this corresponds to a blood flow rate of 5.5 μL/s, or 330 μL/min, which is only slightly higher than the rate estimated in ***fig. 6d***.

## Discussion and Conclusions

In summary, we developed a new method for robust fluorescence enumeration of extremely rare circulating cells directly *in vivo*. In combination with fluorescent dyes, fluorescent proteins and antibody targeted fluorophores^5^, we expect that DiFC will have applications in many preclinical studies involving rare cell types. For example, DiFC can be used to detect early-stage CTC dissemination in animal models of metastasis and response to therapies, for which we have a number such studies already in progress. As we have noted, DiFC builds on our previous work in this area^18^, but encompasses a number of major technical advances (instrument design and signal processing algorithm), that in combination yielded significantly improved instrument capabilities. The measured count rate for DiFC is at least 10 times higher than our previous work with the same animal model, cell line and concentration of injected cells^18^. DiFC also determines cell speeds, depths, and direction of travel which was not possible with our previous designs. We also explicitly tested the sampling volume for the first time.

To reiterate, the main advantage of DiFC is detection sensitivity (sampling rate), which makes it particularly useful for studying rare CTCs. Specifically, our calibration studies showed that DiFC sampled 284 μL of blood per minute of arterial blood in nude mice. This was in excellent agreement with the estimated blood flow rate in the ventral caudal artery, obtained with an independent DiFC measurement of arterial flow speed. We also performed secondary validation of our *in vitro* cell-counting method against conventional flow cytometry, which required and enrichment of blood samples by lysing of red blood cells (***Supplemental fig. S2***).

The DiFC blood sampling rate is also about 2-orders of magnitude higher than microscopy-IVFC methods, which have previously been reported in the range of 0.1-3 μL per min^5^. This is a direct consequence of the fact that DiFC uses diffuse light to interrogate large blood vessels that are generally unsuitable for intravital microscopy. The key enabling features of the DiFC design then are, i) efficient light collection, ii) rejection of non-specific auto-fluorescence, for example of auto-fluorescence generated in the optical fibers due to laser light, iii) rejection of false-positive signals with our matching algorithm, and iv) maintaining blood perfusion in the tail.

On the other hand, it is expected that the minimum detectable cell fluorescent labeling^5,10^ is *lower* for microscopy-IVFC than DiFC, since DiFC generates higher non-specific background than confocal-microscopy. Better quantification of this sensitivity is an ongoing area of research in our group. However, we have tested DiFC with a number of ‘*ex-vivo’* fluorescent dyes (e.g. CTFR and Vybrant DiD), as well as cell lines labeled with constitutively expressed fluorescent proteins, and these are readily detectable with the current design. Use of DiFC with targeted molecular probes is another active area of study.

In addition to being a new stand-alone small animal research tool, we envision that DiFC can be used as a complementary technique to liquid biopsy based technologies such as microfluidic cell-capture and high throughput sequencing^24,25^,. Because DiFC can be used continuously and longitudinally, it can uniquely reveal kinetics and frequency of CTC shedding that occur on the order of minutes, hours and day in particular to establish appropriate time-points for drawing and analyzing blood samples and help inform appropriate time-points for drawing and analyzing blood samples.

Finally, in principle DiFC could be used in larger species and potentially even humans, since it uses highly scattered light in an epi-illumination and detection optical configuration. Theoretically, there are many superficial blood vessels that could be probed with DiFC, for example the cephalic vein in the forearm^26^. However, fluorescent labeling of target cell populations would require the use of molecularly-targeted fluorescent probes, such as those in clinical trials for fluorescence guided surgery^27^. Although this is conceivable, challenges in in specific molecular labeling of CTCs in vivo and regulatory hurdles may outweigh the potential benefits of an *in situ* measurement in humans. As such, the main intended use of DiFC in the foreseeable future is as a new pre-clinical small animal research tool.

## Methods and Materials

### DiFC Instrument

The schematic of the DiFC instrument is shown in ***Fig. 1a***. For the red DiFC system, the light source was a 640 nm diode pumped solid state laser (Excelsior-640C-100-CDRH, Spectra Physics, Santa Clara, CA), the output of which was passed through cleanup band-pass (BP-x; Z640/10x, Chroma Technology, Bellows Falls, VT), and the power was adjusted with a variable ND filter (NDC-25C-2M; Thorlabs Inc, Newton, NJ). The output was split into two beams with a beam-splitter (BS; 49-003; Edmund Optics, Barrington, NJ) and then coupled into the source fibers with an integrated lens fiber coupler (FC-x; F230SMA-B, 633nm anti-reflection coating; Thorlabs). The light power at the sample was 20 mW. The DiFC fiber probes were custom made by EMVision LLC (Loxahatchee, Florida). As shown in ***Fig. 1b***, a micromachined 635/20 nm band-pass filter (BP-f) was mounted over the central source fiber on the probe tip, and a 650 nm long-pass filter (LP-f) ring was mounted over the collection fibers. These drastically reduced autofluorescence generation in the fibers. An aspheric lens (Asph) also improved light coupling. Fluorescent light was collected with an array of 8, 300 μm core multi-mode fibers, which were split into two bundles of 4 fibers. The outputs of the bundles were collimated with integrated fiber couplers (FC-m), filtered with a 700/50 nm band-pass filter (BP-m, ET 700/50m; Chroma), focused with a 25 mm focal length lens (L-m; Edmund) onto the surface of a photomultipler tube (PMT; H6780-20, Hamamatsu, New Jersey). For the experiments here, the band-pass filters (fluorescence detection) were identical, but could be different in the future, for example to allow 2-fluorophore measurements. PMTs were powered by a power supply (C10709, Hamamatsu) The current output of each PMT was amplified with low-noise current pre-amplifiers (PA; SR570, Stanford Research Systems, Sunnyvale, CA), and then digitized with a multi-function data acquisition board (USB-6212 BNC; National Instruments, Austin, TX).

### Data Acquisition and Signal Processing

DiFC data analysis software was specially coded in Matlab and worked as follows (***figure 2c,d***):

- *Step 1. DiFC Data Acquisition:* Data was continuously collected at either 1000 samples per second for arbitrarily long periods of time, typically from 30 to 90 minutes. We summed the measured signal on the two PMTs corresponding to each fiber bundle prior to processing.
- *Step 2. Pre-Processing:* The background autofluorescence was estimated by applying a 2.5 s median filter and then a 7-point moving average filter to the original data traces. The ~10 μA background was subtracted from the raw data, and we then applied a 3 ms moving average filter. We also normalized the amplitude between channels 1 and 2, to partially correct for minor differences in coupling efficiency on the skin surface.
- *Step 3. Identify Cell Candidates:* The software searched for ‘cell candidates’ on the two channels (channel 1, 2) using the Matlab ‘findpeaks’ function and a fixed detection threshold, which was determined using the methodology described below. The minimum peak prominence was 50% of the threshold.
- *Step 4. Analyze Candidate Properties:* The software generated a list of the detection times (***t_1,n_, t_2,m_***), amplitudes (***A_1,n_, A_2,m_***), and estimated cell speeds (***v_c1,n_, v_c2,m_***) of the **N** cell candidates in channel 1 and **M** cell candidates in channel 2. The cell speeds (***v_c1_*** and ***v_c2_***) were estimated by dividing the 1.1 mm detector FOV by the FWHM spike width.
- *Step 5. Identify Candidate Pairs:* The algorithm searched for pairs of ‘similar’ peak-candidates between the two channels in the forward and reverse directions. For each candidate found in channel 1, a search for candidates was performed in channel 2 in an interval corresponding to (***t_1,n_*** + ***d/v_max_***) to (***t_1,n_*** + 10***v_c1,n_*** x ***d)***, where ***d*** was the physical separation between the two collection fibers (3mm), and ***v_max_*** was the assumed maximum possible cell speed of 400 mm/s. Match candidates in channel 2 were then compared to the channel 1 candidate with respect amplitude and speed (***A_2,m_, v_c2,m_***) and a third estimate of the speed ***v_c3_*** = ***d*** / (***t_2,m_*** – ***t_1,n_***). A ‘match’ was defined as a candidate in the interval where amplitudes and speeds agreed within a factor 2.5 on average or less. In the case of multiple matches in the interval, the closest match was selected. In the case where no match was found, the channel 1 candidate was discarded. Coincident peaks (where peak candidates occurred at the exactly the same time on both channels) were also discarded as potential movement artifacts. This analysis was then repeated in the reverse (channel 2-to-1) direction. In rare cases where cells were matched in both forward in reverse directions, the closer match was selected.
- *Step 6. Output:* The analysis reported the count rate in channels 1 and 2 (candidates), and the matched forward (channel 1-to-2) and reverse (channel 2-to-1) directions. It also compiled the speeds, widths and amplitudes of matched spikes.

### Monte Carlo Simulations

We used an open source, GPU-accelerated Monte Carlo program (Monte Carlo eXtreme) to compute the detection sensitivity functions for DiFC (***fig. 5g***)^23^. We modeled the tail as a homogenous 4 mm diameter, 4 cm long cylinder, with voxel size of 250 μm^3^. We used literature values^28^ for optical properties including scattering coefficient (*μ_s_*), absorption coefficient (*μ_a_*), at the excitation (*ex*) and emission (*em*) wavelengths, and the anisotropy coefficient (g) as follows: *μ_s-ex_*= 22 mm^−1^, *μ_s-em_* = 20 mm^−1^ *μ_a-ex_* = 0.002 mm^−1^, μ_a-em_ = 0.0015 mm^−1^, *g* = 0.9.

### MM Cells CTFR Fluorescence Labeling

We used MM.1S multiple myeloma cells (MM.1S.GFP.Luc) that were originally described by Dr. Rosen at Northwestern University. Cells were authenticated by an external service for this study (Bio-Synthesis Inc., Lewisville, TX). MM.1S cells we suspended at 2 x 10^6^ cells/mL in PBS. Labeling solution from CTFR (Cat. C34564) proliferation kit (Thermofisher) were added to a final concentration of 2 μM and incubated for 30 min at 37° C according to the manufacturer’s instruction. After the labeling period, 5 times the original staining volume of phenol free RPMI 1640 with 2 % FBS were added. Cells were centrifuged at 1000 rpm for 5 minutes and then re-suspended in 2 x 10^6^ cells/ml in phenol free RPMI 1640 with 10% FBS. Cells were incubated for an additional 30 min at 37°C. 10ml of phenol free RPMI 1640 with 10% FBS was then added to remove any free dye. Last, cells were centrifuged and re-suspended at the desired concentration for injection.

### Animal Experiments

All mice were handled in accordance with Northeastern University’s Institutional Animal Care and Use Committee (IACUC) policies on animal care. Animal experiments were carried out under Northeastern University IACUC protocol #15-0728R. All experiments and methods were performed with approval from, and in accordance with relevant guidelines and regulations of Northeastern University IACUC.

Female athymic NCr-nu/nu nude mice aged approximately 10 weeks (Charles River Labs, Wilmington, MA) were injected intravenously (i.v.) via the tail vein with 200 μL of either i) 1 x 10^5^ CTFR-labeled plus 10^6^ un-labeled MM.1S cells, or ii) 2.5 x 10^4^ CTFR-labeled plus 10^6^ un-labeled MM.1S cells (N = 5 mice each). In both cases, approximately 10^6^ cells were injected, but in the second case a lower fraction were labeled. The rationale was to produce similar clearance kinetics, since these are known to be dependent on the concentration of the injected cells. Moreover, we note that most injected cells were cleared rapidly from circulation or lost at the site of injection, so that only a small fraction reached quasi-stable circulation.

Mice were scanned with the DiFC system, 10 minutes after injection. Mice were held under inhaled isoflurane throughout the experiments. After approximately 90 minutes we stopped DiFC and drew between 0.5-1 mL of blood via cardiac puncture. Mice were then immediately euthanized. Fluorescently-labeled cells in the blood were counted as below. We also performed sham control injections with 200 μL of PBS (N = 7 mice), which were scanned with DiFC for 60 minutes each.

### Calculation of DiFC Detection Threshold

We used a cell counting threshold of 250 nA for DiFC measurements involving CTFR labeled cells. We first tested the brightness of CTFR-labeled MM.1S cells on DiFC by running them through a bare strand of microbore Tygon tubing (TGY-010-C, Small Parts, Inc., Seattle, Washington) at a concentration of 10^3^ / mL. Samples were placed in a 1 mL syringe mounted in a micro-syringe pump (70-2209, Harvard Apparatus, Holliston, Massachusetts) configured to produce a flow speed of 30 μL/min. We normalized these measurements to the mean brightness of Flash Red 4 microspheres (FR4; Bangs Laboratories, Inc., Fishers, IN) which were measured on the same day. Since FR4 is a stable reference bead this normalization accounted for minor inter-experimental drifts in instrument sensitivity. An example 2-minute trace on CTFR-MM cells in culture is shown in ***Supplemental Figure S1a***, and a histogram of measured brightness for CTFR-MM cells for all experiments is shown in ***fig. S1b***. CTFR-MM brightness was on average 4.4 times greater than FR4 microspheres. We also verified this using a commercial flow cytometer (FC) (Attune NxT, Thermo Fisher Scientific, Waltham, MA) as shown in ***fig. S1c***, which shows the histogram distribution for CTFR-MM cells and FR4 beads. Based on these data, we set a conservative threshold for CTFR-labeled MM.1S cells on either system of 50% of the intensity of FR4 (dotted lines, ***figs. S1b,c***)

The corresponding *in vivo* threshold was then determined by injecting 10^5^ CellSorting microspheres (CS; Cat. C16507; Life Technologies, Carlsbad, CA) in an additional N = 5 nude (nu/nu) mice. CS microspheres were on average 23 times brighter than FR4 spheres. CS microspheres were measured rather than FR4 microspheres since they circulate *in vivo*, whereas FR4 microspheres immediately cleared from circulation. The average measured DiFC signal from CS microspheres *in vivo* was 11.3 μA, so that 50% of FR4 is 1/46 of the measured signal, equivalent to 246 nA *in vivo*. This was rounded to 250 nA.

### Counting of MM-CTFR Cells in Blood Samples

After each DiFC measurement we drew approximately 0.5-1.0 mL of blood and immediately diluted it in 100 μL of 100 units/mL heparin (H3393-50KU, Sigma Aldrich Corp, Natick, MA) mixed with 500 μL of PBS. Diluted blood samples were run through the DiFC in a bare strand of Tygon tubing, followed by suspension of 10^3^ FR4 microspheres per mL of PBS. Fluorescence measurements from cell blood samples were then normalized to the mean intensity of FR4 beads as above. Example data traces from two mice are shown in ***figs. S1d,e***. Peaks above 50% of the mean intensity of FR4 (dotted red line) were counted as a CTFR-labeled MM cell. Smaller peaks were assumed to be debris or cell fragments. This threshold was chosen for consistency with the DiFC counting threshold above, i.e. so that *in vivo* and blood sample measurements could be compared. The number of peaks was divided by the blood volume to estimate the concentration of labeled MM.1S cells in the blood. For example, for the data in ***fig. S1d***, 65 cells were counted in 840 μL of blood = 77.4 cells / mL. For the data in ***fig. S1e***, 15 cells were counted in 590 μL of blood = 25.4 cells/mL.

The rationale for this method is that blood samples could be immediately analyzed with our system with minimal processing. We verified that this method was accurate by spiking whole blood samples with known quantities of MM-CTFR cells and counting them with our system. After counting with DiFC, the same samples were removed and enriched by lysing RBCs with a lysis buffer (420301, Biolegend, San Diego, CA) according to the manufacturer’s instructions. The labeled cells for the suspension were then counted on a flow cytometer (Attune NxT, Thermo Fisher). This was repeated in triplicate (N = 3) at concentrations of 0, 100, 250, 500, and 750 cells/mL. DiFC and FC data are shown in ***Supplemental figure S2a***, showing good agreement between the methods.

Representative flow cytometry plots for this analysis are shown in ***fig S2b-g***. We first ran a solution of FR4 beads (***fig. S2b***) and stock suspension of labeled MM.1S cells (***fig. S2c***) to determine the side scatter (SSC), forward scatter (FSC), and fluorescence distribution of the cells. Spiked blood samples were then analyzed, and an identical SSC and FSC gate applied as the MM.1S CTFR stock suspension, as shown in ***fig. S2d***. This subset of cells was then analyzed for red RL2 fluorescence (710/50 nm filter) as follows: the fluorescence threshold was selected as 50% of the mode intensity of FR4 beads measured from the stock FR4 solutions with the same instrument settings on the same day (***fig. S2e***). Application of this threshold on the SSC-FSC gated blood sample data (***fig. S2f***) removed most of the non-fluorescent blood particles, and the remainder cells in the RL2+ gate were counted (***fig. S2g***).

## Acknowledgements

This work was funded by the National Institutes of Health (R01HL124315; NHLBI). We thank Eric Marple of EMVision LLC for helpful advice and discussion related to the fiber probe design. We thank Prof. Qianqian Fang, (Northeastern University) for assistance using the MCX Monte Carlo software package. We thank Prof. Anne Joutel (INCERM, Paris) for helpful discussion of the mouse vascular anatomy. We also thank Ms. Rebecca Sung and Mr. Zihang Fang for generating the 3D instrument diagrams.

## Additional Information

### Availability of Materials and Data

The datasets generated and analyzed during the current study, as well as the analysis code (written in Matlab) are available from the corresponding author on reasonable request.

### Competing Interests Statement

The authors declare that they have no competing interests.

### Author Contributions

XT and PB developed the DiFC instrument. XT, RP, PB and JR carried out experiments, analyzed data and prepared figures. CL helped plan experiments and supervise the project. MN supervised the project and wrote the manuscript text with support from XT, RP and PB. All authors reviewed and approved the manuscript.

**Supplemental Figure S1.**
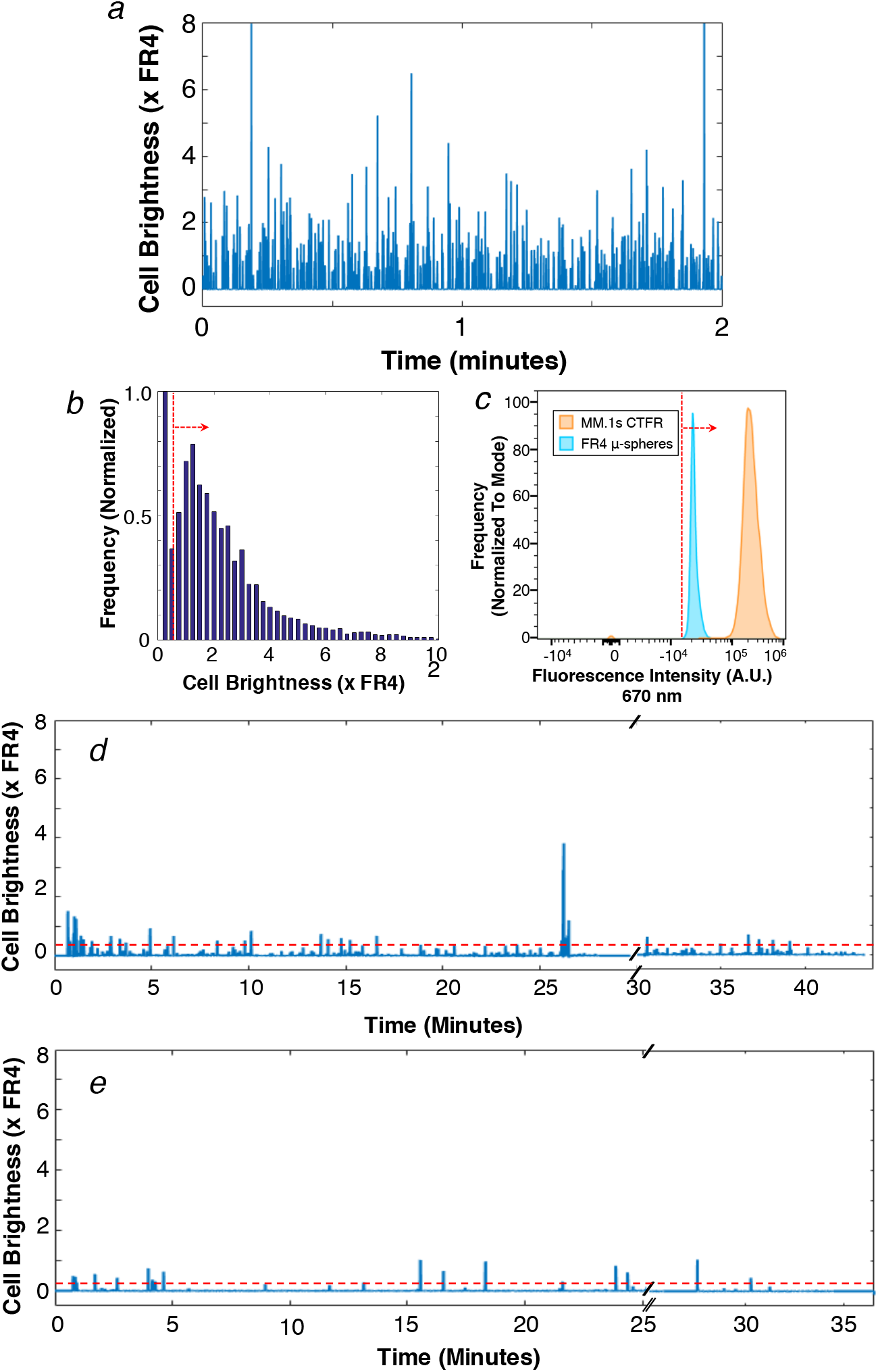
(a) An example DiFC trace of CTFR-labeled MM cells in vitro, normalized to FR4 reference microspheres. (b) Histogram of peak intensities of CTFR- labeled MM cells measured with DiFC, and (c) measured with a flow cytometer. (d,e) Example DiFC data measured from two mouse blood samples. The dotted red lines indicate the counting threshold of 50% of FR4 microspheres. See text for details

**Supplemental Figure S2.**
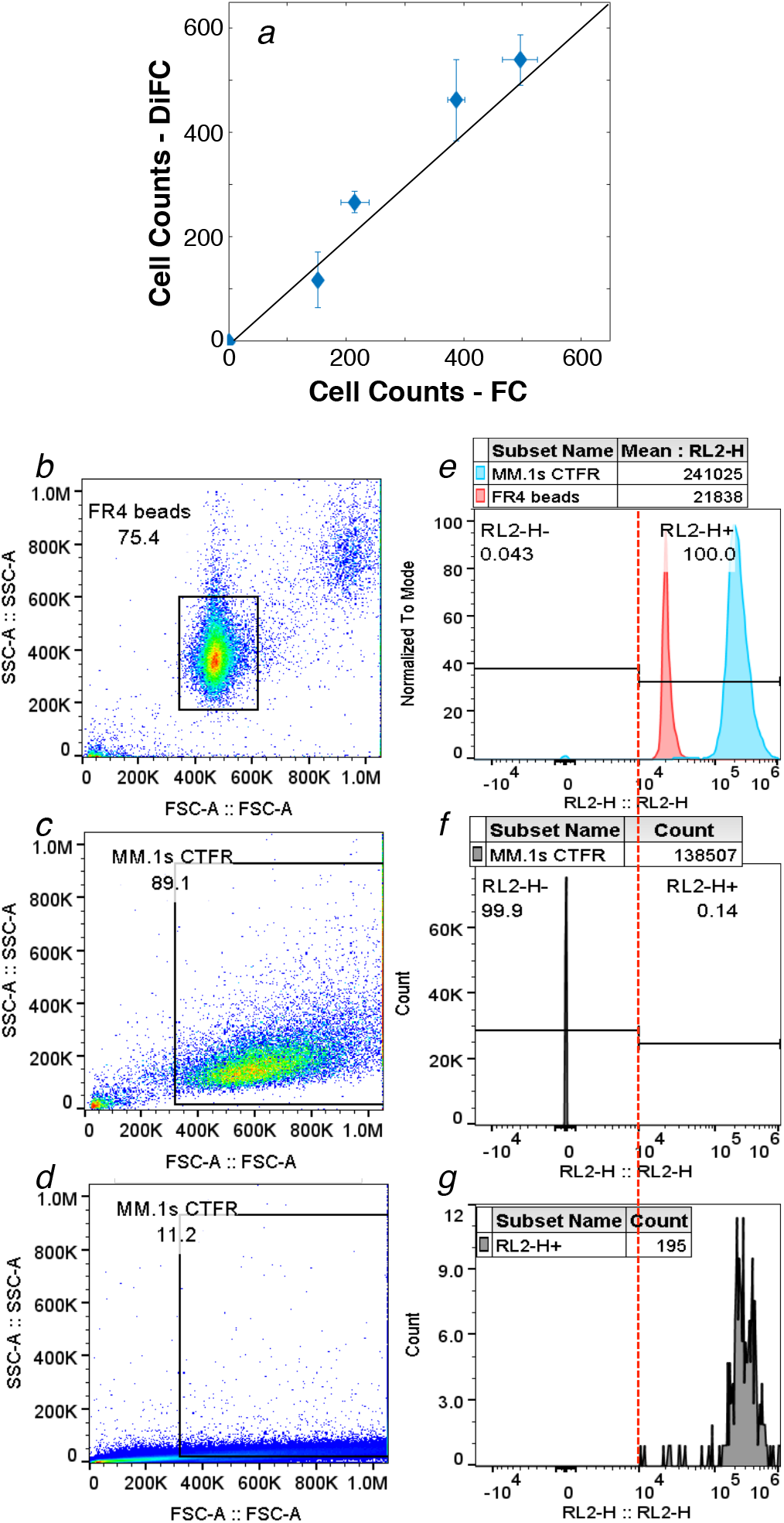
(a) We verified the counting accuracy of DiFC against flow cytometry (FC) for blood samples spiked with CTFR-labeled MM cells. Range bars represent the variability over 3 trials. The FC gating methodology for CTFR is shown: (b) SSC-FSC plot for FR4 reference beads. (c) SSC-FSC plot for CTFR-MM stock cell suspension. (d) SSC-FSC plot for whole blood spiked with approximately 250 CTFR-MM cells. The gate was selected from the CTFR-MM stock suspension in panel (c). (e) Fluorescence (RL2-H, 710/50nm) histogram for FR4 reference beads and CTFR-MM stock suspension, used to select the counting threshold (vertical dotted red line). (f) Fluorescence histogram from the spiked blood sample in the SSC-FSC gate from panel (d). (g) Fluorescence histogram of RL2-H+ cells in the gate.

